# Both the transcriptional activator, Bcd, and transcriptional repressor, Cic, form small mobile oligomeric clusters in early fly embryo nuclei

**DOI:** 10.1101/2024.01.30.578077

**Authors:** Lili Zhang, Lydia Hodgins, Shariful Sakib, Ahmad Mahmood, Carmina Perez-Romero, Robert A. Marmion, Nathalie Dostatni, Cécile Fradin

**Affiliations:** Department of Physics and Astronomy, McMaster University, 1280 Main St. W, Hamilton, ON, L8S 4M1, Canada; Department of Biochemistry and Biomedical Sciences, McMaster University, 1280 Main St. W, Hamilton, ON, L8S 4K1, Canada; The Lewis-Sigler Institute for Integrative Genomics, Princeton University, Princeton, New Jersey, 08544, USA; Institut Curie, PSL University, CNRS, Sorbonne University, Nuclear Dynamics, 75005 Paris, France

## Abstract

Transcription factors play an essential role in pattern formation during early embryo development, generating a strikingly fast and precise transcriptional response that results in sharp gene expression boundaries. To characterize the steps leading up to transcription, we performed a side-by-side comparison of the nuclear dynamics of two morphogens, a transcriptional activator, Bicoid (Bcd), and a transcriptional repressor, Capicua (Cic), both involved in body patterning along the anterior-posterior axis of the early *Drosophila* embryo. We used a combination of fluorescence recovery after photobleaching, fluorescence correlation spectroscopy, and single particle tracking to access a wide range of dynamical timescales. Despite their opposite effects on gene transcription, we find that Bcd and Cic have very similar nuclear dynamics, characterized by the co-existence of a freely diffusing monomer population with a number of oligomeric clusters, which range from low stoichiometry and high mobility clusters to larger, DNA-bound hubs. Our observations are consistent with the inclusion of both Bcd and Cic into transcriptional hubs or condensates, while putting constraints on the mechanism by which these form. These results fit in with the recent proposal that many transcription factors might share a common search strategy for target genes regulatory regions that makes use of their large unstructured regions, and may eventually help explain how the transcriptional response they elicit can be at the same time so fast and so precise.

**SIGNIFICANCE:** By conducting a comparative study of the nuclear dynamics of Bicoid (a transcriptional activator) and Capicua (a transcriptional repressor) in the *Drosophila* embryo, we have uncovered a striking similarity in their behaviours. Despite their divergent roles in transcription, both proteins have a propensity to form oligomeric species ranging from highly mobile, low stoichiometry clusters to larger, DNA-bound hubs. Such findings impose new constraints on the existing models of gene regulation by transcription factors, particularly in aspects related to target search and oligomeric binding to gene regulatory regions needed to explain the rapid and precise transcriptional response observed in developmental processes.

## INTRODUCTION

Transcription factors (TF) play a crucial role in organism development by regulating gene expression in space and time. TFs come in two opposite flavours, activator or repressor, yet they all act on transcription via tight binding to specific sequences in their target genes’ regulatory regions. The interplay between gene activation and repression is especially important in embryos, as it determines the boundaries between gene expression domains eventually leading to tissue differentiation. The position and sharpness of gene expression boundaries is under the tight control of morphogens, molecules that form concentration or activity gradients and affect transcription in a concentration or activity-dependent manner.

Bicoid (Bcd) and Capicua (Cic) are two examples of morphogens that directly act on transcription as they are also TFs. Both are maternally expressed proteins involved in the control of body patterning along the anterior-posterior axis in the early *Drosophila* embryo during nuclear cycles (nc) 8 to 14 (1). Bcd is a homeobox transcriptional activator that forms an exponential concentration gradient (2, 3) and specifies the embryo’s head region (4, 5). It has at least 66 target genes including *hunchback* (*hb*) (6, 7). Cic is a high-mobility group (HMG)-box transcriptional repressor (8). In fly embryos, it is phosphorylated and degraded at both poles due to signalling relayed by the receptor tyrosine kinase Torso (9, 10). It thus represses transcription of its target genes, e.g. *tailless* (*tll*), in the central region of the embryo, and acts as a specificator of terminal regions (8). An example of the entanglement between Bcd and Cic gene regulation is that some genes, like *tll*, are direct targets of both morphogens.

Both Bcd and Cic elicit a remarkably fast and precise transcriptional response. Gene expression in response to Bcd nuclear import is established in only a couple of minutes (11, 12). Cic gene repression triggered by optogenetic methods is complete within the same time frame (13, 14). This points to a very efficient search mechanism to locate target genes. At the same time, small changes in concentration along the embryo anterior-posterior axis are enough to turn on or off the transcription of their target genes, leading to well-defined expression domains (15–18). Cooperative binding to target gene regulatory regions, which often possess multiple binding sites for the same TF, likely drives the sharpness of this response. In the case of Bcd, there is experimental evidence that cooperativity is involved in binding to DNA target sites (7, 19), and it might in part explain the steepness of the *hb* transcriptional response (20–23). Cooperative binding is also thought to be implicated in Cic repression (14). It is unclear, however, how search for target gene and cooperativity play out at the molecular level and affect transcription.

In recent years, evidence has been mounting that molecules participating in transcription assemble into small regulatory hubs (24–29). Some especially clear evidence for the existence of these hubs were found in early fly embryo development (30– 33). Many TFs contain low-complexity sequence domains (LCDs) predicted to form intrinsically disordered regions (IDRs) (34, 35), which may in some cases promote liquid-liquid phase separation (36). Taking this into account, an emerging model of gene expression regulation postulates the existence of transcriptional condensates with elevated concentration of TFs (37–42). By either increasing or buffering the local concentration of TFs at target gene regulatory regions, they could contribute to the speed of the transcriptional response and promote cooperativity (43). Long-lived (on the order of seconds) regions with high Bcd concentration have been observed in the nuclei of early *Drosophila* embryos, using lightsheet imaging and single molecule tracking (30, 31). Yet the mechanism leading to this clustering is unknown.

In this paper, we directly compare the nuclear distribution and dynamics of both Bcd and Cic in the nuclei of nc 13-14 *D. melanogaster* embryos, to check for differences and similarities in their behavior before they specifically bind to one of their target DNA sequences. Using a combination of fluorescence recovery after photobleaching (FRAP), fluorescence correlation spectroscopy (FCS) and single particle tracking (SPT), we were able to cover a wide range of dynamical time scales (0.01 ms ∼100 s) and length scales (300 nm ∼10 *μ*m), and could test in particular for the presence of small and mobile protein clusters. We show that both Bcd and Cic form or associate with protein clusters, ranging from low stoichiometry mobile clusters visible in FCS experiments, up to larger DNA-bound TF hubs visible in SPT experiments.

## RESULTS

### IDRs flank the DNA-binding domains of Bcd and Cic

As transcription factors, both Bcd and Cic have DNA-binding regions, whose structures have been experimentally studied. Bcd has a single homeodomain comprised of three alpha helices (45). Cic, on the other hand, has two DNA-binding regions, a HMG-box complemented by a second DNA binding domain called C1 (the structures of human Cic DNA binding regions have recently been solved (46)). Since no experimental data is available for the other regions of these proteins, we tested the hypothesis that they might contain disordered regions using different IDR predictors, including Spot-dis (47), AUCpreD (48), MetaDisorder (49) and MobiDB-lite (50). In addition, the AI system AlphaFold (which predicts the 3D structure of proteins from their primary amino acid sequence) was used to infer disorder based on the pLDDT score, a per-residue confidence measure. A pLDDT score less than 50 corresponds to very low confidence and is reported to correlate with intrinsic disorder (51). A comparison of the predicted disordered regions reported by these 5 bioinformatic tools for both Bcd and Cic is shown in Fig. 1A. Despite some expected differences in the exact predictions made by these different approaches, large IDRs are predicted by all of them for both proteins on either sides of the central helical DNA binding domains (homeodomain or HMG-box). as illustrated by their AlphaFold structures (Fig. 1B).

**Figure 1.**
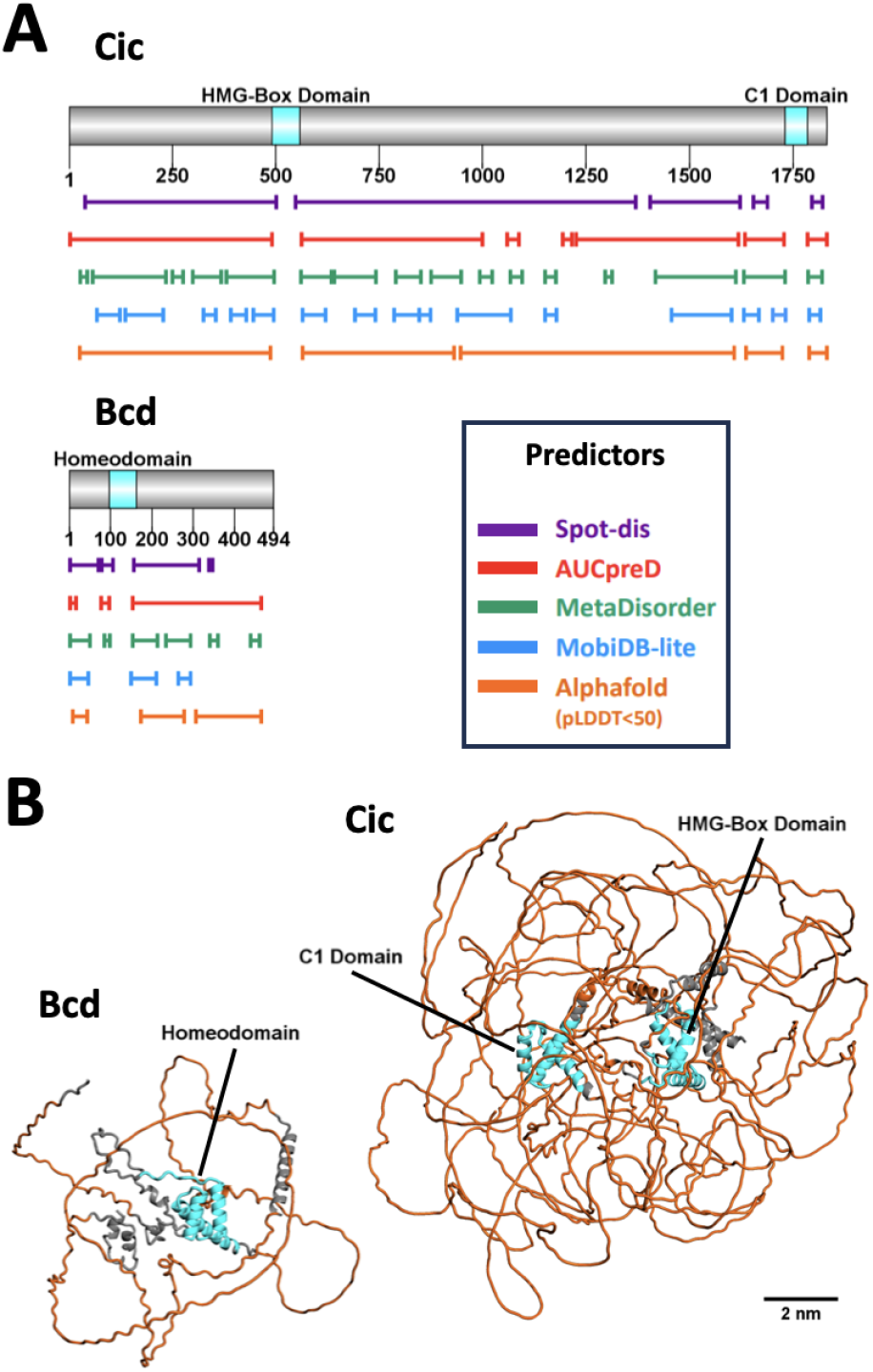
Predicted structures for *D. melanogaster* Bcd and Cic. (A) Predicted IDRs for Cic and Bcd using different bioinformatic tools, rendered using the protein domain illustrator DOG 2.0 (44). (B) Structure predicted by AlphaFold for Bcd (AF-P09081-F1) and Cic (AF-Q9U1H0-F1). Ordered helical DNA-binding regions are shown in cyan in both the amino acid sequence bar and the AlphaFold structures.

### A vast majority of nuclear Bcd and Cic proteins are mobile

To evaluate the large-scale mobility of Bcd and Cic inside nuclei and detect the eventual presence of an immobile fraction, we performed FRAP experiments in *D. melanogaster* embryos expressing either Cic-sfGFP, Bcd-eEGFP or the control NLS-eGFP protein, where half a nucleus (in the mid-embryo cortical region, at nc 13 or 14) was photobleached and the fluorescence recovery monitored for 20 s afterwards. All experiments were conducted in the region of the embryo situated mid-way between the anterior and posterior poles, where the nuclear concentrations of Bcd and Cic are comparable.

It can be immediately inferred from the recovery curves shown in Fig. 2A-C that none of the three studied proteins has a significant immobile fraction on the timescale of the experiments, as they all homogeneously redistribute across the nucleus in a few seconds. For Bcd-eGFP and NLS-eGFP (Fig. 2B,C), the initial fast intranuclear redistribution is followed by a slower overall increase in total nuclear fluorescence due to nucleo-cytoplasmic exchange (which occurs on a timescale of ≃1 min as shown by whole nucleus FRAP experiments, see Supplementary Material, Fig. S1, and Ref. (52)). In contrast, no overall nuclear fluorescence recovery is observed for Cic (Fig. 2A), whose nucleo-cytoplasmic transport is severely restricted (with an average nuclear residence time > 20 min in the middle region of embryos, see Supplementary Material, Fig. S1, and Ref. (10)). Recovery curves were fitted with an exponential function accounting for intranuclear redistribution complemented, in the case of Bcd-eGFP and NLS-eGFP, by a linear term accounting for the slow net influx of cytoplasmic proteins (Eq. 1). This allowed extracting the characteristic intranuclear recovery time (τ_*f*_) as well as the fraction of the signal contributed by immobile proteins. This analysis shows that all three proteins have an immobile fraction of less than 5% (Fig. 2D).

**Figure 2.**
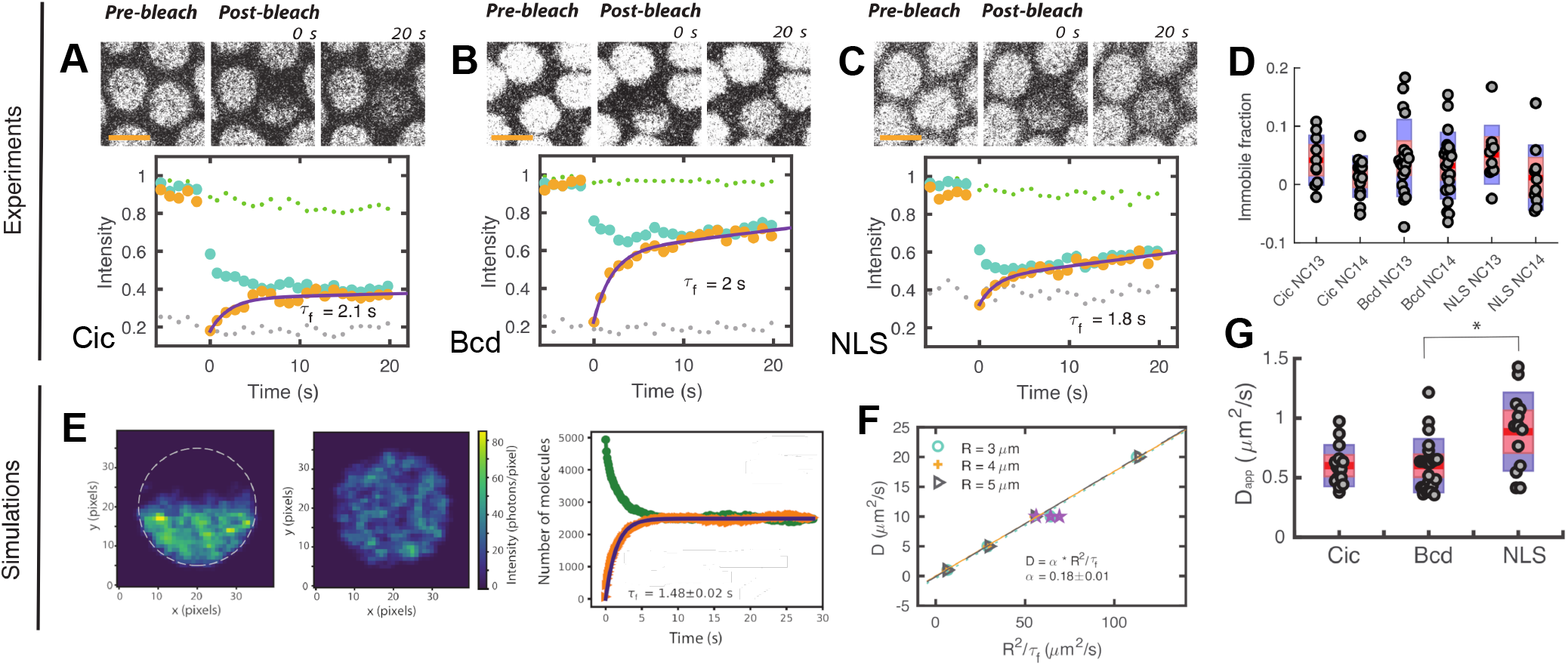
Nuclear mobility of Bcd and Cic captured by FRAP. (**A-C**) Representative intranuclear FRAP experiments in embryos expressing Cic-sfGFP (**A**), Bcd-eGFP (**B**), or NLS-eGFP (**C**). Top: Confocal images of the nuclei pre-photobleaching, and at *t* = 0 and 20 s post-photobleaching (scale bar: 5 *μm*). Bottom: Recovery curves for the intact (large green symbols) and photobleached (large orange symbols, fitted with Eq. 1) halves of the nucleus, along with the average fluorescence of surrounding nuclei (small green symbols) and average background fluorescence between nuclei (small grey symbols). (**D**) Immobile fractions obtained from comparing the recovery curves from the lower and upper halves of nuclei. (**E**) Representative simulation of an intranuclear FRAP experiment, performed for 1000 fluorescent molecules diffusing with *D* = 1 *μ*m^2^/s in a *R* = 3 *μ*m radius nucleus. Simulated confocal images of the system are shown at time t = 1.6 s and t = 60 s after photobleaching, along with the recovery curves for the lower (green symbols) and upper (orange symbols, fit with an exponential function) halves of the nucleus. (**F**) Relationship between *D* and *R*^2^ /τ_*f*_ obtained for simulations performed for different values of *R* and *D*. Star symbols indicate simulations performed for photobleached regions either slightly smaller or slightly larger than half of the nucleus. The line is a linear fit of the data, which returned a proportionality coefficient *α* = 0.18. (**G**) Values of the apparent diffusion coefficient measured in different nuclei for the three studied proteins. The mean is indicated by a red line, the 95% confidence interval by a red box, and the standard deviation by a purple box. The symbol (*\**) indicates a t-test *p*-value < 0.05.

Extracting diffusion coefficients from FRAP experiments requires taking into account the geometry of the photobleached region and, in the case of a finite compartment such as the nucleus, boundary conditions (53, 54). To this end, we performed Monte Carlo simulations to obtain an empirical relationship between the protein diffusion coefficient (*D*), the nucleus radius (*R*) and the observed fluorescence recovery time (τ_*f*_). In the simulation, post-bleach conditions were reproduced by randomly placing fluorescent particles in the lower half of a sphere representing the nucleus, then allowing them to diffuse within that sphere (see Materials and Methods for details). The simulated fluorescence signals in the lower and upper halves of the nucleus (Fig. 2E) reproduces experimental observations, with the exception of the nucleo-cytoplasmic contribution which was not simulated in this case. The characteristic recovery time due to intranuclear diffusion (τ_*f*_) is obtained by fitting the simulated recovery curve for the upper half of the sphere with a single exponential function. From dimensional analysis we expect *D* = *αR*^2^ /τ_*f*_. Our simulations (in which *R* was varied from 3 to 5 *μ*m and *D* from 1 to 20 *μ*m^2^/s) confirmed this relationship and showed that *α* = 0.18 ±0.01 (Fig. 2F). An apparent diffusion coefficient *D*_app_ = *αR*^2^ /τ_*f*_ can then be calculated for each protein (Fig. 2G). However, for all three proteins, τ_*f*_ is only slightly longer than the 1 s photobleaching step, indicating that protein redistribution has already started when the first data point post-photobleaching is acquired. Thus *D*_app_ only represents a lower limit for the protein diffusion coefficient (55, 56). In conclusion, for all three proteins, the overwhelming majority (> 95%) of the nuclear pool is fully mobile, with a motion consistent with diffusion and *D* > 0.6 *μ*m^2^/s for Cic-sfGFP and Bcd-eGFP, and *D* > 0.9 *μ*m^2^/s for NLS-eGFP.

### Bcd and Cic form mobile oligomeric clusters

For a finer comparison of the dynamics of Bcd and Cic, and to closely examine the different mobile states of these proteins, we turned to FCS. Previous FCS studies have established that Bcd has several dynamic modes in nuclei, as evidenced by the presence of at least two separate decays in the collected autocorrelation functions (ACF) (16, 57), and we recently reported a similar behavior for Cic (14). In order to find out wether the slower of these two decays might correspond to an oligomeric species, we developed a novel approach, where series of up to 20 consecutive single-point FCS experiments were performed in a single nucleus while using a relatively high (20 *μ*W) excitation intensity. This allowed observing the evolution of the relative amplitude of these two components under photobleaching conditions, which we show below depends on the stoichiometry of the observed species.

Measurements were performed at the centre of nuclei in the middle region of nc 13 or 14 embryos expressing either Cic-sfGFP, Bcd-eGFP or NLS-eGFP, with representative ACFs shown in Fig. 3A-C. The decay corresponding to protein mobility was observed around 1 *−* 100 ms, between that attributed to photophysics (in the *μs* range) and that due to the progressive photobleaching of the nuclear pool of fluorescent proteins (in the *s* range). The decay due to continuous photobleaching is prominent in the first ACF of each series (orange symbols in Fig. 3A-C), then progressively disappears in subsequent measurements as the fluorescence signal stabilizes. As previously reported (14, 16, 57), the ACFs recorded for Bcd and Cic have a mobility decay term reflecting a range of dynamic processes. To estimate decay times and relative amplitudes associated with these dynamic processes while avoiding over-interpreting the data, ACFs were fitted assuming two diffusing populations of particles (two-component model, Eq. 4, and continuous lines in Fig. 3A-C). The photobleaching decay was taken into account by adding a dedicated term to the ACF (see section and Ref. (58)), which allowed extracting dynamic information from the very first curves obtained in each series, when the signal-to-noise ratio is the highest, and before photobleaching has started significantly affecting the brightness of eventual oligomers. From these fits, four characteristic times are obtained: the photophysics relaxation time τ_*T*_ *≈*20 *μ*s, the characteristic diffusion times of the fast and slow diffusing fractions τ_Df_ *≈*1 *m*s and τ_Ds_ *≈*10 - 100 *m*s, and the photobleaching relaxation time τ_*P*_ *≈*40 s. The measured values of these four parameters, remarkably constant over time, are shown for all three proteins in Fig. 3D-F. Four series of measurements (each lasting either 5, 10, or 20 s) are shown for each protein, demonstrating the consistency of these results.

**Figure 3.**
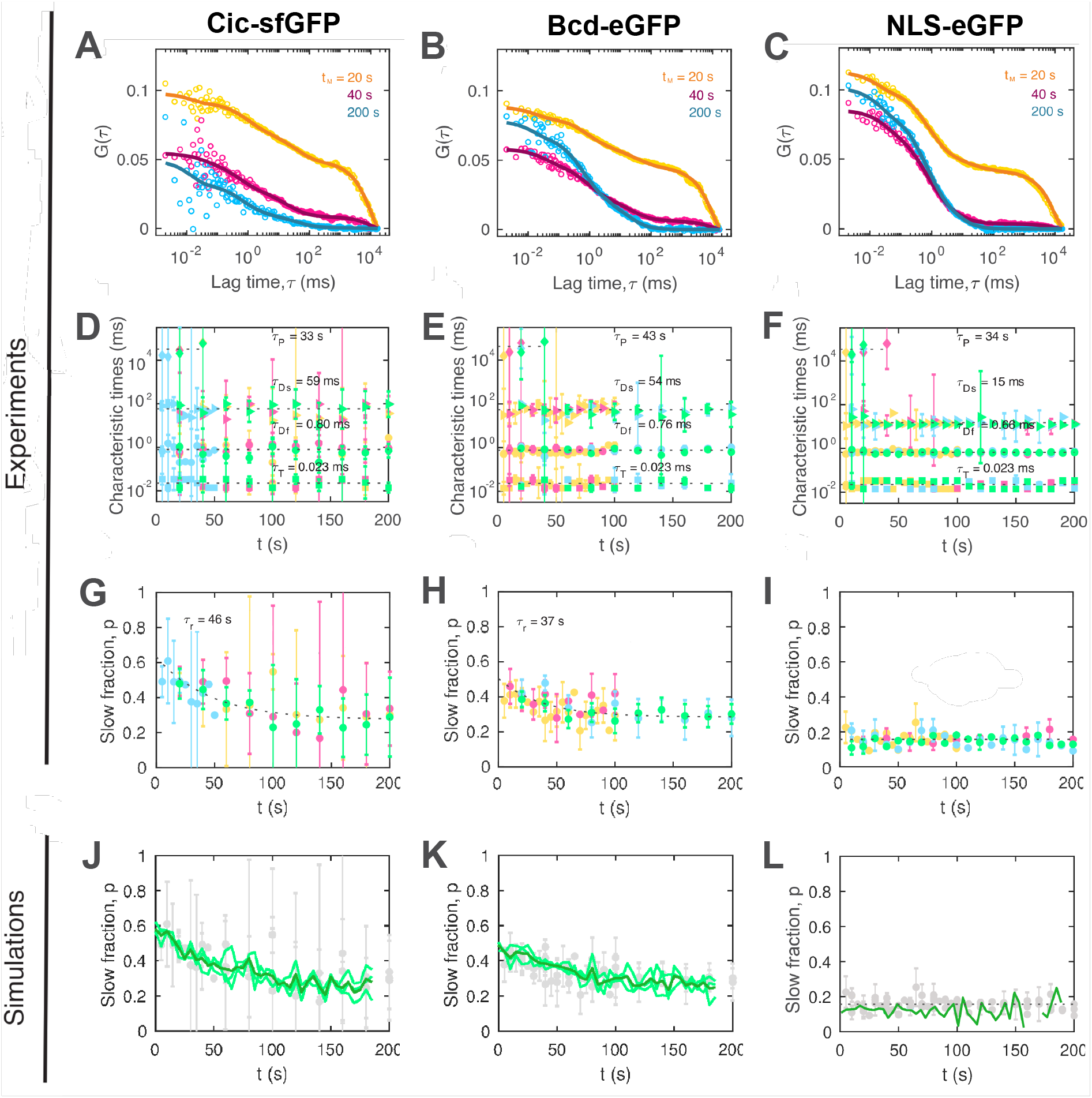
Nuclear mobility of Bcd and Cic characterized by FCS. (**A-C**) ACFs obtained from the first (orange), second (pink) and tenth (blue) measurements in series of 20 consecutive 20 s FCS measurements performed in nuclei of embryos expressing either Cic-sfGFP (**A**), Bcd-eGFP (**B**), or NLS-sfGFP (**C**). Lines are fit with a two-component model taking into account photobleaching. (**D-F**) Characteristic times obtained from the fit of the ACFs: photophysics relaxation time, τ_*T*_, fast diffusion characteristic time, τ_Df_, slow diffusion characteristic time, τ_Ds_, and photobleaching time, τ_*p*_, for Cic-sfGFP (**D**), Bcd-eGFP (**E**) and NLS-eGFP (**F**). The value of τ_*p*_ is obtained only for the first couple of measurements in each series, when the photobleaching contribution to the ACF is substantial. (**G-I**) Relative amplitude of the slow dynamics term, *p*, shown for Cic-sfGFP (**G**), Bcd-eGFP (**H**) and NLS-eGFP (**I**). Different colors represent separate series of FCS measurements. The error bars are 50% confidence interval. (**J-L**) Simulation of the evolution of *p*, when a nucleus containing *N* = 10, 000 fluorophores is submitted to photobleaching (green lines, average of three independent repeats), and compared to the data already shown in (**G-I**) (grey symbols). For **J**,**K** fluorophores were assumed to exist both in a monomeric form (*N* _*f*_ monomers, diffusion coefficient *D* _*f*_ = 15 *μm*^2^/*s*) and in oligomeric clusters (*N*_*s*_ clusters, diffusion coefficient *D*_*s*_ = 0.4 *μm*^2^/*s*). Each clusters initially contained *n* fluorophores, with *n* drawn from a Poisson distribution. For Cic-sfGFP *N*_*s*_/*N* _*f*_ = 0.04 on average *n* = 7 (**J**), while for Bcd-eGFP *N*_*s*_/*N* _*f*_ = 0.08 and on average *n* = 3.5 (**K**). For **L** fluorophores were assumed to alternate between a diffusing state and an immobile, bound state (stick-and-diffuse model) with en equilibrium constant *k*_off_/*k*_on_ = 7.

The mobility of all three proteins is dominated by the fast component (τ_Df_ ≃ 0.8 ms, corresponding to a fast diffusing species with apparent *D* _*f*_ ≃ 30 *μm*^2^ /*s*). In the case of Bcd-eGFP and Cic-sfGFP, this fast component coexists with a significantly slower component (τ_Ds_ ≃ 50 to 60 ms) that could correspond to a larger species with diffusion coefficient *D*_*s*_ ≃ 0.4 *μm*^2^ /*s*. The relative contribution of this slow component (*p*) exponentially decreases over the course of a continuous series of FCS experiments, from ≃ 50 to 25 %, with a characteristic time τ_*r*_ ≈ *≈*40 s (Fig. 3G,H). For the control protein NLS-eGFP, a second dynamic component is also observed, however it is rather fast (τ_Ds_ = 15 ms) and its relative contribution is both small (≃ 10 %) and constant over time (Fig. 3I). The fact that τ_*r*_ is very similar to the photobleaching relaxation time τ_*P*_ suggests that the decrease in *p* observed for Bcd-eGFP and Cic-sfGFP is linked to the gradual photobleaching of the nuclear population of fluorophores. This decrease can be explained if we assume that fast particles contain a single eGFP molecule whereas slow particles contain *n* eGFP molecules. In that case, photobleaching will affect both types of fluorescent particles very differently: for the fast particles, the number of still fluorescent particles (*N* _*f*_) will progressively decrease but not their brightness (*B* _*f*_, always equal to that of a single eGFP molecule), while for the slow particles the brightness of individual particles (initially *B*_*s*_ = *nB* _*f*_) will decrease in a stepwise fashion, while initially their number remains constant. Since for two diffusing species, 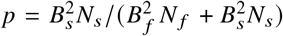 (Eq 5), a decrease in *B*_*s*_ dominates over a decreases in *N* _*f*_ and causes a decrease in *p*. Making a few simplifying assumptions allows to relate the initial and final values of *p* to the quantities *N*_*s*_ /*N* _*f*_ and *n* = *B*_*s*_ /*B* _*f*_, as captured in Eqs. 9 (see Materials and Methods). This allows a first rough estimate of the stoichiometry of these clusters: *n* = 5.0 for Cic and *n* = 2.7 for Bcd.

For a more precise estimate of *n*, one needs to take into account effects that are not captured in Eqs. 9, such as the uneven distribution of fluorescent species within the nucleus due to continuous photobleaching. To that end, we performed simulations of nuclei containing two types of diffusing fluorescent particles, a fast monomeric species with brightness *B* _*f*_, and a slower oligomeric species with average brightness *B*_*s*_ = *nB* _*f*_ and a Poisson distributed number of fluorescent monomers (see Materials and Methods for details). The diffusion coefficients of these particles were chosen to match the values measured for Cic-sfGFP and Bcd-eGFP: *D* _*f*_ = 15 to 30 *μ*m^2^ /s for the monomers and *D*_*s*_ = 0.4 *μ*m^2^/ s for the oligomers. At each time step, particles were subjected to photobleaching with a probability proportional to the local intensity of a laser beam focused at the center of the nucleus. The continuous photobleaching resulted in a progressive decrease of the number and brightness of the different fluorescent particles, and in a progressive decrease of *p* (calculated using Eq 6 for the particles present at the laser focus to simulate the result of an actual FCS experiment). To reproduce the observed *p* (*t*), we varied the simulation parameters and found that we had to use values of *n* that were a slightly higher than predicted by Eqs. 9 (Fig. 3J,K). Simulations were also performed assuming a stick-and-diffuse model (59, 60), where monomers binds and unbinds from DNA, meaning that the slow species corresponds to a transiently immobilized monomer (see Supplementary Material). In that case no variation in *p* was observed (Fig. 3L), as expected because of the fast exchange between the two types of particles. In conclusion, whereas transient binding of monomers can explain the data obtained for the control NLS-eGFP protein, in the case of Bcd-eGFP and Cic-sfGFP the variation of the relative amplitude of the slow dynamic term of the ACF is instead consistent with the presence of slowly diffusing oligomeric clusters with *n* 7 for Cic-sfGFP and *n* 3.5 for Bcd-eGFP. It does not preclude these clusters from transiently binding to DNA.

### Confocal imaging reveals the presence of both mobile and immobile Bcd and Cic clusters

If Bcd and Cic form slowly diffusing clusters with a molecular brightness larger than that of a single GFP molecule, it should be possible to image and track them. To confirm the result of our FCS experiments, we thus turned to fast confocal imaging. Ventral mid section nuclei in nc 13 or 14 embryos expressing each of the three different proteins of interest were acquired at a relatively high frame rate (2 fps) for a total of 20 frames. For both Cic-sfGFP and Bcd-eGFP there appeared small diffraction-limited regions of higher intensity which persisted at the same position over multiple frames, such that when averaging the pixel intensities over 20 frames these regions became more clearly visible (Fig. 4A,B). In contrast, for NLS-eGFP, although regions of higher intensity did occasionally appear in some frames, they did not persist, and were not apparent in average images (Fig. 4C). The FIJI-Mosaic plugin was used to track higher intensity particles and generate distributions of step sizes (see Materials and Methods for details). On average, 3 such particles were detected in nuclei of embryos expressing fluorescent Bcd or Cic, and less than 1 for those expressing NLS-eGFP (Fig. 4D), and the length of the detected trajectories was longer for Cic-sfGFP and Bcd-eGFP than for NLS-eGFP (Fig. 4E). The step size distributions for Cic-sfGFP or Bcd-eGFP (Fig. 4A,B) suggested the presence of two different types of clusters with distinct mobilities (in contrast to what was observed for NLS-eGFP or for a sample of immobilized fluorescent beads, see Fig. 4C). They were thus fit with a two-component Rayleigh distribution, returning in each case two separate diffusion coefficients, the first *≈*0.01 *μ*m^2^/ and the second *≈* 0.05 *−* 0.1 *μ*m^2^/ e.

**Figure 4.**
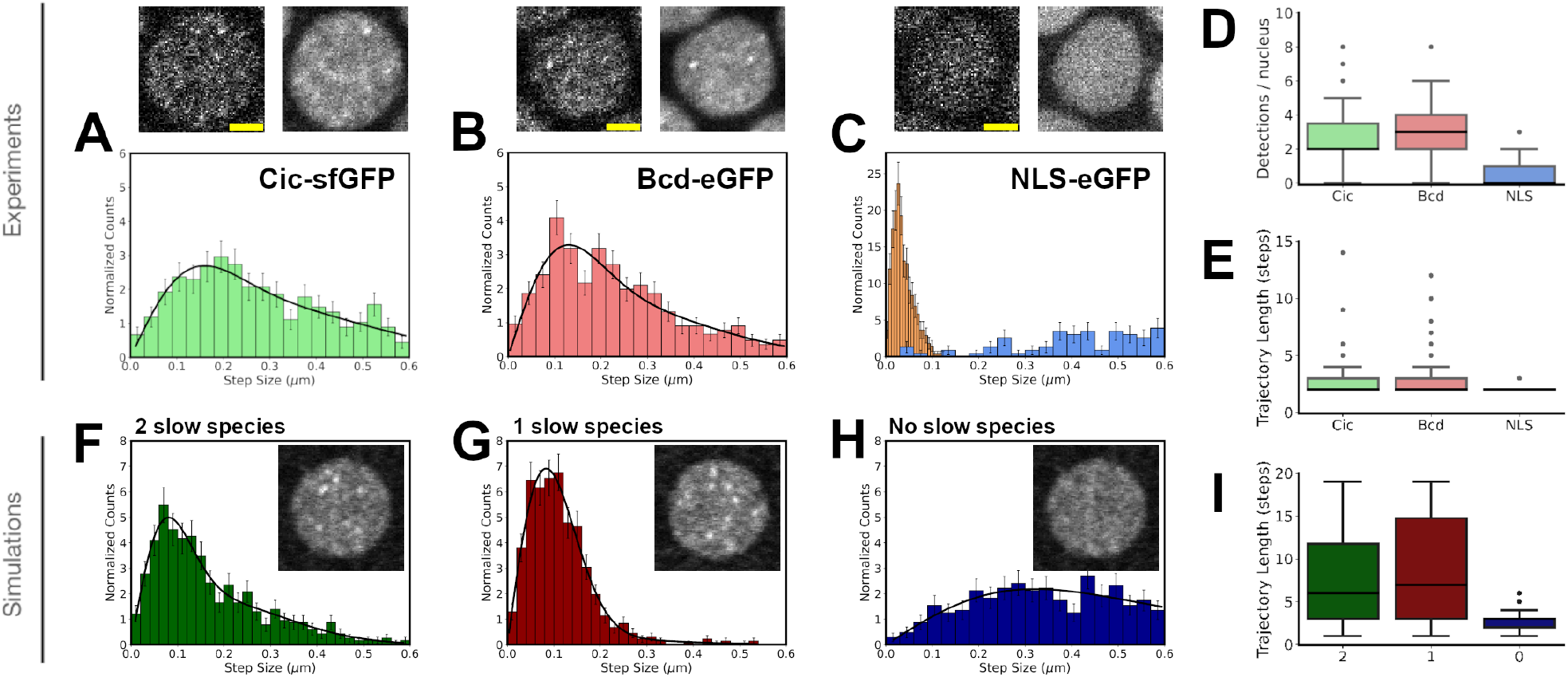
Fluorescent cluster detection in confocal images. (A-C) Confocal images of nuclei acquired in embryos expressing Cic-sfGFP (A), Bcd-eGFP (B) or NLS-eGFP (C). In each case, the first image in a series of 20 (frame rate: 2 s^*−*1^) is shown (top left), along with the average image for that series (top right). The histograms below show the step size distributions (Δ*t* = 0.5 s) for the particle trajectories detected in each type of embryo (for 63 to 67 separate nuclei in each case). For the NLS-eGFP control (C), the step size distribution recorded for immobilized beads (orange) is shown alongside the step size obtained for the protein (blue, from 63 separate nuclei). For Cic-sfGFP and Bcd-eGFP, distributions are fitted with a two-component Rayleigh distribution (Eq. 10). Scale bar is 2 *μ*m. (D) Number of particles detected per nucleus and (E) distribution of trajectory lengths for each type of embryo. (F-H) Step size distributions (Δ*t* = 0.118 s) for particles detected in simulations of nuclei containing, in addition to the fast diffusing monomer population (*D* _*f*_ = 30 *μ*m^2^ /s), either two (F), one (G) or no (H) population(s) of slow bright particles. Each distribution was constructed from analysis of 5 different simulated nuclei. Examples of simulated nuclei images are shown as an inset. (I) Trajectory lengths simulations with two, one or no slow species.

For comparison, we simulated confocal movies of nuclei containing diffusing fluorescent proteins (*D* _*f*_ = 30 *μ*m^2^ /s), to which were added one or two slow species (*D*_*s*_ = 0.2 *μ*m^2^ /s and *D*_*i*_ = 0.015 *μ*m^2^ /s, diffusion coefficients were chosen at the upper limit of our experimental observations) with higher brightness (*B*_*s*_ = 12*B* _*f*_ to simulate particles containing 12 GFP). Photobleaching was not considered since we were simulating images acquired over a short 10 s period with a scanning laser beam. The same analysis procedure was used on these movies to track visible high-intensity particles as was used on the experimental movies. For simulated images including one or two slow species, higher brightness clusters could clearly be seen by eye, and the step size distributions obtained after tracking these clusters resembled a one- or a two-component Rayleigh distribution as expected (Fig. 4F,G). The diffusion coefficients obtained from these fits were close to those of the simulated particles: 0.02 to 0.03 *μ*m^2^/ s, a bit larger than the actual diffusion coefficient of the slowest simulated species likely due to position detection accuracy, and 0.15 *μ*m^2^ /s, a bit lower than the actual diffusion coefficient of the second slow simulated species likely due to steps larger than 0.6 *μ*m not being detected. The faster particles were both too fast and too dim to be detected, just like the fast fraction of the proteins was not detected in the experimental movies. The step size distribution obtained for the system containing 2 types of slow particles closely resembles what was observed in experiments for Bcd and Cic (Fig. 4F). For the simulations that included only fast particles, higher intensity regions could also sometimes be observed in individual images, however, when these regions were tracked the step size distribution was noticeable broader than for the other simulations (Fig. 4H) and the trajectory lengths were much shorter (Fig. 4I), closely resembling what was observed for NLS-eGFP.

## DISCUSSION

The aim of this study was to directly compare the intranuclear dynamics of Bcd and Cic, focussing on their role as TFs, as previous studies had centered instead on one protein or the other, often in the context of gradient formation (10, 52, 56, 57, 61). We were expecting similarities due to their shared role as TF in the early fly embryo (with a need for fast action due to rapid nuclear divisions) and comparable structures (Fig. 1), as well as differences due to their ultimately opposite functions (activator vs. repressor). Overall, we find that the intranuclear mobilities of Bcd and Cic are surprisingly alike (as summarized in Table 1), with both proteins found in three different forms: *(1)* abundant freely diffusing monomers, *(2)* slowly diffusing oligomeric clusters, and *(3)* rare immobile oligomeric hubs (as illustrated in Fig. 5).

**Table 1:** Summary of the physical parameters measured in this study for the different observed species for each of the three studied proteins. The superscript at the end of the parameter name refers to the method used to measure it - a: nuclear FRAP, b: intranuclear FRAP, c: FCS followed by two-component model analysis of ACF, d: analysis of repeated FCS experiments under photobleaching conditions e: single particle tracking.

| Species | Parameter | Symbol | Cic | Bcd | NLS | Unit |
| --- | --- | --- | --- | --- | --- | --- |
| Monomers | Nucleo-cytoplasmic exchange<sup>a</sup> | $\tau_b$ | $> 20$ | $1.3 \pm 0.4$ | $1.3 \pm 0.2$ | min |
| | Diffusion coefficient<sup>b</sup> | $D_f$ | $> 0.6 \pm 0.2$ | $> 0.6 \pm 0.2$ | $> 0.9 \pm 0.3$ | $\mu\text{m}^2/\text{s}$ |
| | Diffusion coefficient<sup>c</sup> | $D_f$ | $< 28 \pm 18$ | $< 30 \pm 8$ | $< 34 \pm 5$ | $\mu\text{m}^2/\text{s}$ |
| Mobile clusters | Oligomeric fraction<sup>d</sup> | $N_s/N_f$ | $\approx 0.04$ | $\approx 0.08$ | $\approx 0$ | |
| | Stoichiometry<sup>d</sup> | $n$ | $\approx 7$ | $\approx 3.5$ | | |
| | Diffusion coefficient<sup>c</sup> | $D_s$ | $< 0.38 \pm 0.20$ | $< 0.42 \pm 0.19$ | | $\mu\text{m}^2/\text{s}$ |
| | Diffusion coefficient<sup>e</sup> | $D_s$ | $> 0.082 \pm 0.028$ | $> 0.056 \pm 0.013$ | | $\mu\text{m}^2/\text{s}$ |
| Immobile hubs | Total immobile fraction<sup>b</sup> | | $2 \pm 4$ | $4 \pm 6$ | $3 \pm 6$ | % |
| | Oligomer immobile fraction<sup>e</sup> | $N_i/N_s$ | $< 0.51$ | $< 0.76$ | | |
| | Diffusion coefficient<sup>e</sup> | $D_i$ | $0.011 \pm 0.006$ | $0.007 \pm 0.003$ | | $\mu\text{m}^2/\text{s}$ |

**Figure 5.**
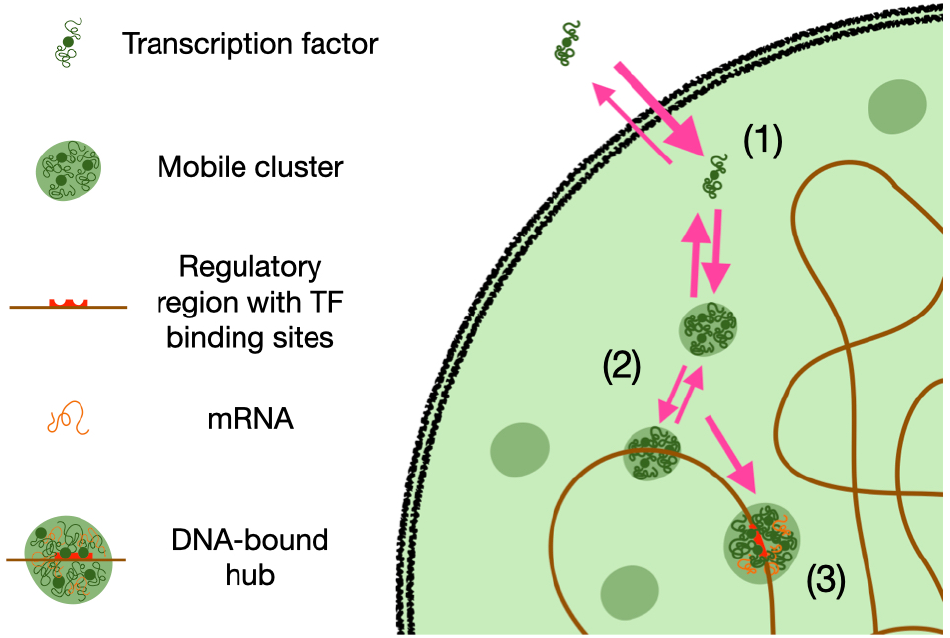
A possible model for TF dynamics based on the nuclear species detected in this study. In nuclei TFs coexist in at least three different forms: (1) freely diffusing monomers in equilibrium with a cytoplasmic pool, that can quickly scan the nuclear space, (2) low stoichiometry slowly diffusing TF clusters, possibly formed *via* low-specificity protein-protein interactions mediated by their large unstructured regions - these clusters might include a number of different proteins and might also be loosely interacting with nuclear structures and (4) larger immobile transcriptional hubs specifically bound to promoter regions and likely including mRNA transcripts.

### Existence of an intermediate between freely diffusing TFs and transcriptional hubs

Most current models of TFs nuclear dynamics propose that a fraction of these proteins are freely diffusing monomers (or homodimers), making occasional non-specific contacts with the DNA until they find the promoter region of a target gene, where they might join or lay the foundations for a transcriptional hub (here we use the term “hub” to designate a protein-rich cluster bound to DNA) (37, 62–65). These models are based on long-standing observations of a fast dynamics mode for TFs, accompanied by the more recent realization that TFs form DNA-bound foci or hubs (24–29).

A freely diffusing form is indeed observed in all FCS nuclear studies of either Bcd or Cic (it is too fast to be captured by most SPT and FRAP experiments), including this one, where it manifests as a distinct decay around ∼1 ms in the ACF (Fig. 3). Here we used a slightly higher excitation intensity than in previous works, in order to cause continuous photo-bleaching. The values we extract for the diffusion coefficient of the Bcd-eGFP and Cic-sfGFP monomers (*D* _*f*_ *≈* 30 *μ*m^2^ /s) are thus biased towards higher values, and should be considered as upper bounds. Actual values are probably closer to those previously reported for Bcd, i.e. *D* _*f*_ = 8 *−*12 *μ*m^2^/ s (16, 57). Still, this abundant supply of diffusing TFs (50% of the total nuclear pool) is remarkably fast, and able to quickly explore the nucleus in search of binding partners.

The existence of DNA-bound Bcd hubs at transcription sites is well established (30, 31). In this work, immobile hubs are observed in time-lapse confocal images, not only for Bcd, but also for Cic., and shown to have a very small apparent diffusion coefficients, *D*_*i*_ *≈*0.01 *μ*m^2^/ *s*, within the range of diffusion coefficients reported for chromatin elements *in vivo* (31, 66, 67). (Since chromatin diffusion is restricted, meaningful apparent diffusion coefficient comparisons can only be made considering measurements performed on the same *≈*1 s timescale as ours). These hubs persist for at least a few seconds, a dynamics consistent with TF specific binding to a target DNA sequence. They represent at most a few % of the total Bcd or Cic pool, as expected if they are indeed organized around the regulatory regions of target genes (both Bcd and Cic have tens of target genes on the *Drosophila* genome (68, 69), still a small number compared to thousands of Bcd and Cic molecules present in each mid-embryo nucleus).

One unexpected aspect of our study is that small mobile oligomeric clusters are detected for both Bcd and Cic, and visible in both FCS and SPT experiments. In FCS experiments, they appear as a distinct decay in the ACFs around 50 ms (Fig. 3D,E). This decay has been reported in every FCS study of either Bcd (16, 56, 57) or Cic (14). For Bcd, it has also been shown to shift to faster timescales upon impairment of Bcd DNA-binding ability (57). The novelty of our study lays in our analysis of the slow decrease of the amplitude of this term under continuous photobleaching conditions, which allows us to conclude that it belongs to slowly diffusing oligomeric clusters containing on average ≃ 3.5 Bcd or ≃ 7 Cic molecules, and persistent on the *≈*30 s continuous photobleaching characteristic timescale. These clusters diffuse with an apparent diffusion coefficient between ∼0.05 (FCS) and ∼0.5 *μ*m^2^ /s (SPT), suggesting a range of sizes, with FCS more sensitive to smaller faster clusters, and SPT to larger slower ones. With a diffusion coefficient about 100-fold smaller than that measured for the free protein, their average hydrodynamic radius could be as large as a few hundred nanometers (i.e. 100-fold larger than a monomer). This relatively large size would still be consistent with their diffraction-limited appearance in confocal images, and with a low Bcd or Cic copy number if their bulk is made up of scaffolding proteins such as maternal Zelda (known to support Bcd hub formation (31)). However, the clusters are likely to be quite a bit smaller than that, and their low mobility due at least in part to transient interactions with chromatin.

### Role of Bcd and Cic clusters in transcription

As a number of TFs, including at least one repressor (70), have been observed to form noticeable hubs (24–27, 29–31), many current transcription models have immobile TF hubs play an important role in transcription regulation, e.g. by increasing the local concentration of TFs in the vicinity of target genes, by pulling together different target genes or by buffering the concentration of free TFs (37, 40, 43, 64). But what of the relatively abundant smaller mobile clusters observed in this study? We speculate that there are several ways in which they may assist transcription. First, they could be a vehicle for TFs to find their target genes via facilitated diffusion, alternating between three-dimensional diffusion and short one-dimensional searches along the DNA. While facilitated diffusion of isolated TFs can in principle significantly speed up target search (71, 72), a lone protein would find any other protein decorating the DNA to be a formidable obstacle. Instead, a small oligomeric structure containing several TFs (and therefore several DNA-binding domains), would be able to jump over obstacles. A larger “searcher” might also have a larger non-specific affinity for DNA and facilitate translocation between adjacent DNA segments (27). Finally, facilitated diffusion of oligomeric clusters brought together via IDR interactions would be consistent with the observation that IDRs increase the efficiency with which TFs recognize their target sequences (73, 74). Second, TF cluster formation might explain the observed cooperativity of the binding of Bcd (19) and Cic (14) to DNA, which was shown to be mediated, in the case of Bcd, by its intrinsically disordered regions (19)). Small clusters might help pre-package important factors for transcription in a small mobile particle still uncommitted to a particular target gene, and delivery of this package would promote binding of not just one, but several TF to the same region. Third, the existence of two distinct pools of mobile TFs (monomers and oligomeric clusters) might explain why the activity gradient of Bcd differs from its concentration gradient (12) and how Bcd is capable of both concentration-dependent and concentration-independent transcription activation (68). If mobile TF clusters play a role in transcriptional activation, then our work suggests they also play a role in transcriptional repression. A common mechanism for these two processes would go a long way in explaining the similarities observed in the dynamics and precision of the transcriptional response elicited by Bcd and Cic. Although our study is focused on the early fly embryo, Cic has a homolog gene in human. Further, since most TFs possess the same mix of DNA-binding and disordered regions as Bcd and Cic, and since a slow fraction is present in most FCS studies of TFs (including p53 (75), the retinoic acid receptor (76), Hox TFs in *Drosophila*, and a number of mammalian embryo TFs (77)) most of the conclusions of this study are not restricted to morphogens or to the fly embryo, and instead likely to be very general. In conclusion, we propose that a comprehensive transcription model should apply to both transcriptional activation and repression, and take into account the presence of small TF oligomeric clusters.

### Cluster formation mechanism

The largely disordered structures of Bcd and Cic suggest the possibility that the clusters we observe are formed by phase separation, since IDRs facilitate the type of multivalent interactions necessary for this process (35, 78). However, IDR on TFs have also been implicated in promoting multivalent interactions without phase separation (79). For Bcd, immobile clusters have been observed in posterior nuclei at vanishingly low Bcd concentration (30), and previous FCS experiments have shown that the slow dynamic component we attribute here to mobile Bcd clusters is still present in posterior nuclei (although less abundant than in anterior nuclei) (57). This is inconsistent with a simple phase transition involving one single player (Bcd), but it is still possible that a phase transition occurs with the help of co-factors (80), for example maternal Zelda which is uniformly expressed throughout the embryo at this stage (81), or nucleic acids since the DNA binding domain of Bcd has been implicated in the formation of these clusters (57). Our own observation that the formation of immobile hubs bound to DNA is accompanied by a significant stable population of small mobile clusters, is also inconsistent with a simple phase transition mechanism. To be explained in the context of liquid-liquid phase separation, they would require invoking cluster size-limiting arguments such as the involvement of an elastic network (82–84).

## MATERIALS AND METHODS

### IDR prediction from protein sequence

As bioinformatic IDR predictors vary in their approaches to predict disorder, we used several of them, mostly chosen according to the performance ranking published by Nielsen *et al*. (85). Namely, we used the top ranked machine-learning predictors Spot-dis (47) and AUCpreD (48), and we used the meta model Metadisorder (49) which combines multiple predictive algorithms to report a final consensus (86). We also used MobiDB-lite (50), another meta model developed after the analysis by Nielsen *et al*. was published, and AlphaFold (51, 87). Spot-dis, AUCpreD, and MetaDisorder provide a per residue disorder probability ranging from 0 to 1. Using the homeodomain of Bcd as a reference for structured residues, we manually set the threshold value for disorder to 0.9 for Spot-dis and AUCpreD, and to 0.7 for MetaDisorder. MobiDB-lite offers binary results in which a residue is either structured or disordered. For AlphaFold, residues with a pLDDT below 50 were considered likely to be disordered and a ±5 buffer was used so that a single isolated residue predicted as ordered or disordered was excluded.

### Experiments

#### Drosophila embryo preparation

*Drosophila* embryos were prepared for imaging following the protocol detailed in ref. (88). Briefly, *D. melanogaster* fly strains expressing NLS-eGFP, Bcd-eGFP (a kind gift from Dr. Wieschaus) (52) or Cic-sfGFP (a kind gift from Dr. Shvartsman) (14) were maintained in a 25°C incubator with alternating day-night lighting. To collect embryos for experiments, the plastic tube containing flies was inverted on a collection plate with yeast paste in the centre. After about 3 hours, the embryos laid on the collection plate were placed using a tweezer on a double-sided tape to manually remove the chorion. Dechorionated embryos were then transferred to a thin line of heptan glue on a 0.17 mm coverslip. The ventral side of the embryo was placed in contact with the coverslip such that as many nuclei as possible in the embryo could be observed in a single field of view just above the surface of the coverslip. Lastly, a small drop of Halocarbon oil 700 (Sigma Aldrich, St. Louis, MI, US) was added on top of the embryos to prevent water evaporation while allowing oxygen permeation for necessary metabolism. All subsequent experiments were performed at room temperature.

#### Confocal imaging

Time series of confocal images of nuclei in the ventral cortical layer of the embryo in contact with the coverslip were acquired with an Ti2-E inverted confocal microscope (Nikon Canada, Mississauga, ON, Canada) using an oil immersion Apo objective (60×, NA 1.4) and high-sensitivity GaAsP photomultipliers tubes. A 488 nm laser excitation and FITC emission bandpass filter (521/42) were used for fluorescence imaging. Acquisition settings for all movies were: laser power: 5% (100 *μ*W), confocal aperture: 1 Airy unit (29.4 *μ*m), pixel dwell time: 4.1 *μ*s, pixel size: 0.1 *μ*m, detector gain: 65, image size: 256 × 256 pxl, number of frames: 20, continuous acquisition mode. The resulting frame rate was 2 fps. When needed, masks corresponding to nuclei in the images were created using a machine-learning based algorithm implemented in the software Ilastik (89).

#### Intranuclear FRAP

FRAP experiments were performed on a Nikon eclipse Ti inverted microscope fitted out with an oil-immersion Nikon Plan Apo *λ* objective (60×, NA 1.4) used in conjunction with a 38 *μ*m confocal aperture. Fluorescence excitation and photobleaching were achieved with a 488 nm laser. A rectangular region covering one half of a nucleus was photobleached for 1 s using a laser power of 80% (0.4 mW). A 128 ×128 pxl region around that nucleus was imaged imaged once before photobleaching and for 20 s after at 1 s intervals using a 5.1 % (25 *μ*W) laser power, a 2.7 *μ*s pixel dwell time and a 0.1 *μ*m/pixel size. The software Fiji was used to extract the intensity of the bleached and unbleached halves of the nucleus (90). Fitting of the recovery curves was done using MATLAB (MathWorks, Natick, MA, US).

#### FCS

Single-point FCS data were acquired with an Insight Cell confocal microscope (Evotec Technologies, Hamburg, Germany, now PerkinElmer, Waltham, MA, USA). Excitation was performed using a 488 nm continuous wave solid state diode-pumped laser (Sapphire 488-20/460-10, Coherent, Santa Clara, CA, USA). The excitation beam was set to overrfill the back-aperture of the water-immersion objective (UApoN, 40×, 1.15 NA, Olympus Canada, Richmond Hill, ON, Canada) and used in conjunction with a 40 *μ*m pinhole. Calibration measurements were performed with solutions of Alexa Fluor 488 (Life Technologies, Carlsbad, CA, USA), which has a known diffusion coefficient *D* = 435 *μ* m^2^/s at 22.5 °*C* (91). The measured characteristic diffusion time of Alexa Fluor 488 on this instrument was τ_*D*_ 55 *μ*s, from which the 1 /*e*^2^ radius of the detection volume 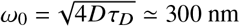 could be calculated. ACFs were analyzed using either FCS+plus Analyze (Evotec) or MATLAB.

## Data analysis

### Intranuclear FRAP

Two separate processes were assumed to contribute to fluorescence recovery in half-photobleached nuclei: 1) the redistribution of fluorescent proteins inside the nucleus due to intranuclear diffusion (characteristic time τ_*f*_), and 2) the slower exchange of molecules between nucleus and cytoplasm due to nucleo-cytoplasmic transport (characteristic time τ_*b*_). Thus, the evolution of the mean fluorescence intensity in the bleached and unbleached halves of the nucleus are given by:

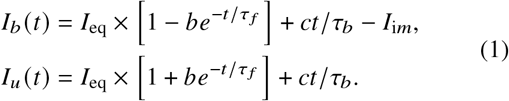

*I*_eq_ is the equilibrium intensity that would be reached in the unbleached half of the nucleus in the absence of nucleo-cytoplasmic transport, *b* and *c* are constants, and *I*_i*m*_ is the mean intensity contributed by immobile fluorescent particles. Note that in the expressions above we assumed that the exchange between eventual different diffusing populations would be rapid enough on the time scale of this experiment that they would appear as a single population. Further, since for these experiments *t* «τ_*b*_ we used a linearized expression for the slow recovery term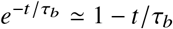.

The fraction of immobile molecules, *p*_im_, can be calculated from the experimentally accessible quantities *I*_*u*_ (*t*), *I*_*b*_ (*t*), and *I*_0_(*t*) (mean intensity of a control unbleached nucleus) for *t* » τ_*f*_ :

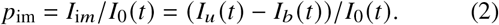

In practice, the mean from the last five data points in each experiment was used to calculate *p*_im_.

### FCS

Following the work done in (58), ACFs obtained as the result of nuclear FCS measurements were analyzed as the sum of two contributions:

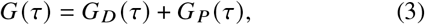

where *G*_*D*_ (τ) accounted for the fluorophore dynamics (diffusion, transient immobilization, photophysics, etc…) and *G* _*P*_ (τ) accounted for the gradual decrease in fluorescence due to continuous photobleaching in the small nuclear space.

#### Two-component model

Protein dynamics was modelled assuming two species of diffusing fluorescent molecules were present (diffusion coefficients 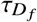and 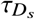). The part of the ACF reflecting fluorophore dynamics then is (92):

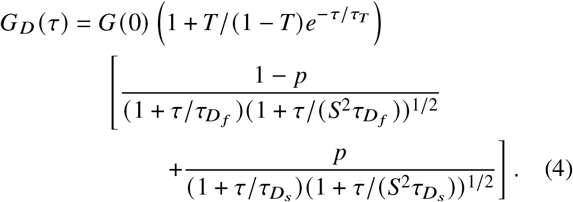

*S* is the aspect ratio of the detection volume. *T* is the fraction of molecules in the dark state with a relaxation time of τ_*T*_. The diffusion coefficients of the fast and slow species are related to characteristic diffusion times via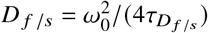. *G*(0) is the amplitude of the diffusive part of the ACF. The relative amplitude of the slow diffusive term is given by (93, 94):

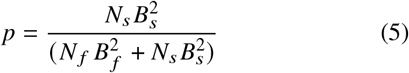

where *N* _*f*_ /_*s*_ and *B* _*f*_ /_*s*_ are the concentration and molecular brightness of the fast and slow particles, respectively. If we consider that the fast species is a monomer whereas slow particles have a distribution of molecular brightness *B*_*s,i*_, each with probability *p*_*i*_, this formula can be generalized to:

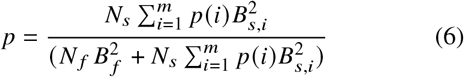

#### Continuous photobleaching term in the ACF

The photo-bleaching contribution is given by (58):

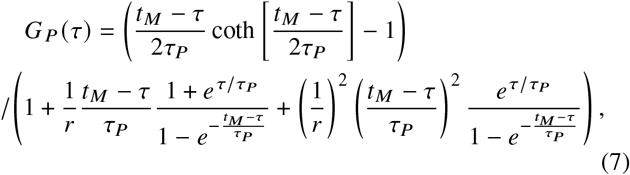

where *t*_*M*_ is the FCS measurement time, τ_*P*_ is the photo-bleaching characteristic time and *r* is the signal-to-noise at the beginning of the measurement (58).

#### Amplitude of the slow diffusive term and photobleaching

To gain an intuition for the effect of continuous photobleaching on the result of FCS experiments done with mixes of fluorescent particles of different stoichiometries, we examined a simple situation. We considered a small compartment containing two distinct non-interchangeable populations of diffusing particles: monomers containing a single fluorophore (brightness *B*) and clusters containing a fixed number *n* of that same fluorophore (brightness *nB*). We further considered that the probability for a fluorophore to photobleach remained the same whether it is part of a monomer or a cluster. Finally we assumed that the concentration of both species remained constant throughout the compartment, i.e. that monomers and clusters are diffusing fast enough to avoid the build up of a photobleaching “hole” at the laser focus (95).

Due to photobleaching, the total fluorescence contributed by each species will decrease more or less exponentially, with a time constant τ_*P*_. For monomers, this means that the concentration of particles that are still fluorescent (initially equal to *N*_*m*,0_) decreases exponentially over time: 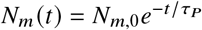. Meanwhile, the brightness of these still-fluorescent monomers remain constant, *B*_*m*_ (*t*) = *B*. In contrast, for clusters, photo-bleaching will reduce their brightness before it reduces their number. In the limit of large clusters, and of small time, we can approximate their brightness as 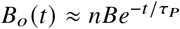, and their concentration as constant, *N*_*o*_ (*t*) *≈N*_*o*,0_. The relative amplitude of the term due to the slower oligomeric clusters (Eq. 5) is then given by:

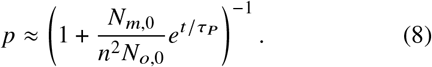

Eq. 8 predicts that, initially, *p* = *p*_0_ = (1 +*N*_*m*,0_ /(*n*^2^*N*_*o*,0_)) ^*−*1^, but that, as photobleaching proceeds, *p* relaxes towards a lower value with a time constant τ_*P*_.

However, when the average brightness of clusters starts approaching *B* (i.e. when *t* ≃ τ_*P*_ ln *n*), the assumption that *N*_*o*_ (*t*) is constant breaks down, and Eq. 8 is no longer valid. Instead, *p* reaches a constant value around *t* τ_*P*_ ln *n*, since at this point clusters only have one fluorescent unit left on average and start acting as monomers from the point of view of photobleaching, with exponentially decreasing concentration and constant brightness. At equilibrium we therefore expect *p ≃ p*_*∞*_ = 1/ (1 +*N*_*m*,0_ /(*nN*_*o*,0_)). An estimate of the relative concentration and stoichiometry of the clusters can thus be obtained from the observed values of *p*_0_ and *p*_*∞*_:

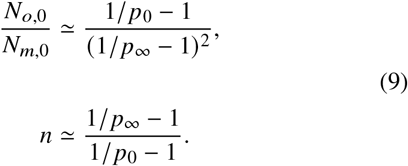

### Particle tracking

The FIJI-Mosaic plugin (90, 96) was used to track high intensity regions within confocal image series (acquired or simulated). For experimental images, a Gaussian blur filter (kernel radius = 0.2 *μ*m) was first applied to reduce the effects of noise. Tracking parameters wiithin the Mosiac program were kept constant. The expect particle radius was set to 4 pxl (0.4 *μ*m, slightly larger than the diffraction limit of the instrument), the cutoff radius to 0.001 and the percentile to 0.4%. The linkrange and maximum allowed displacement between frames were kept at the low values of 1 frame and 6.0 pxl (0.6 *μ*m), respectively, to avoid erroneously linking detection events generated by different particles. Trajectories consisting of only one step were excluded from the analysis. For each condition, a single one-step distribution of displacements was produced from all detected trajectories in all images. The diffusion coefficients of the particles were estimated by fitting this probability density with a two-population Rayleigh probability density function:

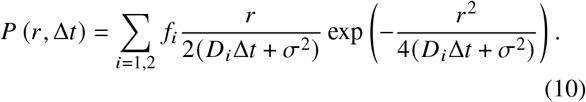

*r* is the length of the step. Δ*t* is the time between two consecutive steps. *f*_*i*_ and *D*_*i*_ are the fraction and diffusion coefficient of population *i*. The localization precision, *σ*^2^, was evaluated to be *σ*^2^ = 3 ×10^*−*3^ *μ*m^2^ in experimental confocal images (from images of immobilized beads), and *σ*^2^ = 1.7 ×10^*−*4^ *μ*m^2^ in simulated confocal images (from simulations of immobilized particles) - see Supplementary Material for details. Analysis of step size distribution was used instead of analysis of mean-squared displacement (MSD), both to better allow distinguishing between different populations and because many of the trajectories were too short for meaningful MSD to be calculated. A Kruskal-Wallis test was applied to both the samples of trajectory lengths and the samples of the number of trajectories detected per nucleus to quantify statistically significant differences between the three proteins.

### Simulations

Different Monte Carlo simulations of the three-dimensional motion of diffraction-limited fluorescent particles inside a single nucleus were performed using code written in Python. A 12 ×12 ×12*μm*^3^ simulation box was used in all simulations. A sphere (with a 3 to 5 *μm*-radius) at the centre of the cubic box represented the nuclear envelope delimiting the nucleus. Initial particle positions were determined either according to thermal equilibrium or to reflect a particular non-equilibrium initial configuration, as created for example by photobleaching. Each particle was defined by its diffusion coefficient, *D*, and by the number of fluorescent proteins it contained, *n*, resulting in a molecular brightness *B* = *nb*, where *B* is the molecular brightness of a single fluorescent protein.

#### TF motion

At each simulation step (each corresponding to a duration *δt* = 1), each particle was allowed to move in each spatial direction by a step size drawn from a Gaussian distribution with a mean of 0 and a variance of 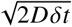. To simulate two-component diffusion, particles were randomly assigned one of two states at creation (with diffusion coefficient *D* _*f*_ for “fast-diffusing” particles and *D*_*s*_ for “slow-diffusing” particles), then allowed to move accordingly throughout the rest of the simulation. Moves bringing particles outside of the simulation box were refused. Moves causing particles to cross the nuclear envelope were accepted with a probability chosen to reflect the probability of import into (*p*_*in*_) or export from (*p*_*out*_) the nucleus. Nuclear transport was always prohibited for particles representing clusters (*p*_*in*_ = *p*_*out*_ = 0).

#### Confocal Imaging

Simulated confocal images were produced at different time points during a simulation for visualization or continuously to produce a movie of the system. In confocal imaging, the laser focus is scanned line by line and point by point across a *n*_*x*_ × *n*_*y*_ rectangular field of view and at each point the signal is recorded in a single photon detector. Similarly in the simulations, the signal was calculated at each pixel sequentially, while allowing molecules to move between pixel recordings. To match experimental conditions, a pixel size (distance between two consecutive pixels) *d* = 0.1 *μ* m/pixel and a pixel dwell time τ = 4.1 *μ*ms or τ = 1 ms were used. For an image started off at time *T*, the *i*th pixel on the *j* th line would be recorded at time *t* = *T* + ((*j −* 1)*n*_*x*_ + *i*)τ, thus the signal contributed by particle *k* to that pixel was calculated as:

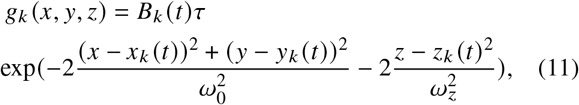

where *x*_*k*_ (*t*), *y*_*k*_ (*t*), *z*_*k*_ (*t*) is the position of particle *k* at time *t*, and *B*_*k*_ (*t*) its molecular brightness (in photon/s). (*x, y, z*) is the position of the pixel, where *x* = *id* and *y* = *j d. ω*_0_, *ω*_*z*_ are the short and long 1/*e*^2^ radii of the Gaussian confocal detection volume, respectively. The average signal at each pixel was then calculated as the sum of the signal contributed by each particle, plus a constant background intensity *I*_bckg_ (97). When needed, the effect of photon noise was taken into account by replacing, at each pixel, the calculated average signal by a value drawn from a Poisson noise with mean and variance equal to that average signal. When simulating time-lapse confocal images, an image size of 120*x*120 pxl and a frame rate of 8.5 fps were used. The code used to simulate time-lapse confocal images is available at github.com.

### Continuous photobleaching

To simulate continuous photobleaching during a single-point FCS experiment, we considered the light intensity profile of a laser (wavelength *λ*) focused at the centre of the nucleus (*x* = 0, *y* = 0, *z* = 0) (98):

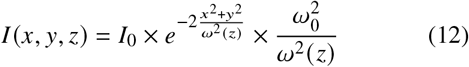

where:

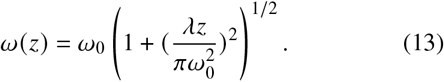

The probability that a fluorophore at position *x*_*k*_, *y*_*k*_, *z*_*k*_ would photobleach during the small time interval *δt* was considered to be *P*_*P*_ (*x*_*k*_, *y*_*k*_, *z*_*k*_) = *a* ×(*I* (*x*_*k*_, *y*_*k*_, *z*_*k*_) / *I*_0_)*δt*, where *a* is a constant which, in an experiment, would depend on the laser intensity and the photostability of the fluorophore. The value *a* = 3 was used in all the simulations presented here. The code used for simulating continuous photobleaching is available at github.com.

## AUTHOR CONTRIBUTIONS

L.Z. designed the research, performed and analyzed all the FRAP and FCS experiments, and wrote the manuscript. L.H. acquired and analyzed the confocal images, performed the bioinformatics study and wrote the manuscript. S.S. performed fly stock control experiments. A.M. developed the image segmentation workflow. C.P.-R. performed preliminary FCS experiments. R.A.M. and N.D. helped design the research. C.F. designed the research, performed analytical calculations, and wrote the manuscript.

## ACKNOWLEDGMENTS

We thank S. Y. Shvartsman for many helpful discussions, S. Rauscher for her help with secondary structure prediction tools and M. Rose for his early contribution to the confocal image simulation software. This research was funded by the Natural Sciences and Engineering Research Council of Canada through a discovery grant to C.F. (RGPIN-2015-06362). It was enabled in part by resources provided by Compute Ontario and the Digital Research Alliance of Canada. Confocal imaging was done at the Centre for Advanced Light Microscopy at McMaster University. L.Z. and C.P.-R. were both recipients of an Ontario Trillium Scholarship. L. H. was supported by an NSERC-USRA scholarship.

## SUPPLEMENTARY MATERIAL

This article has a supplementary information file.

## Notes

### Competing Interest Statement

The authors have declared no competing interest.

