## Supplementary Information for "Both the transcriptional activator, Bcd, and transcriptional repressor, Cic, form small mobile oligomeric clusters in early fly embryo nuclei"

### 1 Nucleo-cytoplasmic exchange

The kinetics of nucleo-cytoplasmic transport for Cic-sfGFP, Bcd-eGFP and NLS-eGFP was studied by performing whole nucleus FRAP experiments. Using a Nikon eclipse Ti inverted microscope (60× NA 0.95 air-immersion Plan Apo $\lambda$ objective, 46 $\mu$m pinhole) a small ∼2 $\mu$m-diameter circular region at the centre of a nucleus was photobleached for 3 s using the full 0.5 mW power of a 488 nm laser. A 1024×1024 pixels region around that nucleus was imaged once before the photobleaching step and for 10 to 20 min after at 30 s intervals, using a lower 30 $\mu$W laser power and a pixel dwell time of 4.6 $\mu$s. While the nuclear fluorescence of Bcd-eGFP and NLS-eGFP mainly recovers after a few minutes, that of Cic-sfGFP does not, even after 20 mins (Fig. S1A).

An automatic segmentation procedure was performed to extract nuclei position using a machine learning algorithm encoded within the software Ilastik (Berg et al., 2019). This allowed computing the total fluorescence intensity recorded from each nucleus in the field of view (Fig. S1B,C). The first 10 min of the recovery curves were fitted with a double exponential function, the first accounting for the fluorescence recovery due to nucleo-cytoplasmic transport (characteristic time $\tau_b$), and the second for continuous photobleaching that occurred while imaging (characteristic time $\tau_p$):

$$I_b(t) = I_\infty[1 - ae^{-t/\tau_b}]e^{-t/\tau_p}, \tag{S1}$$

where $I_\infty$ represents the equilibrium intensity that would be reached in the absence of continuous photobleaching and $a$ is the fraction of initially photobleached molecules. The recovered fraction ($f$) was obtained by comparing the mean fluorescence in the photobleached nucleus ($I_b(t)$) to that in nearby nuclei ($I_u(t)$) at $t = 20$ min post-photobleaching, taking into account the fluorescence background ($I_{bckg}$):

$$f = [I_b(t = 20 \text{ min}) - I_{bckg}] \,/\, [I_u(t = 20 \text{ min}) - I_{bckg}] \,. \tag{S2}$$

The average recovery times, $\tau_b$, and recovery percentages at 20 min, obtained from repeated experiments, are shown in Fig. S1D,E. Assuming that the cytoplasmic concentration of Bcd-eGFP and NLS-eGFP is constant, the recovery time depends only on the proteins' nuclear export rate ($k_{\text{out}}$), where $\tau_b = 1/k_{\text{out}}$. In effect, $\tau_b$ thus corresponds to the average time spent by the protein in the nucleus. The nuclear export rates measured for Bcd-eEGFP and NLS-eEGFP are shown in Fig. S1F). All measured or inferred parameters related to nucleo-cytoplasmic transport are listed in Table S1.

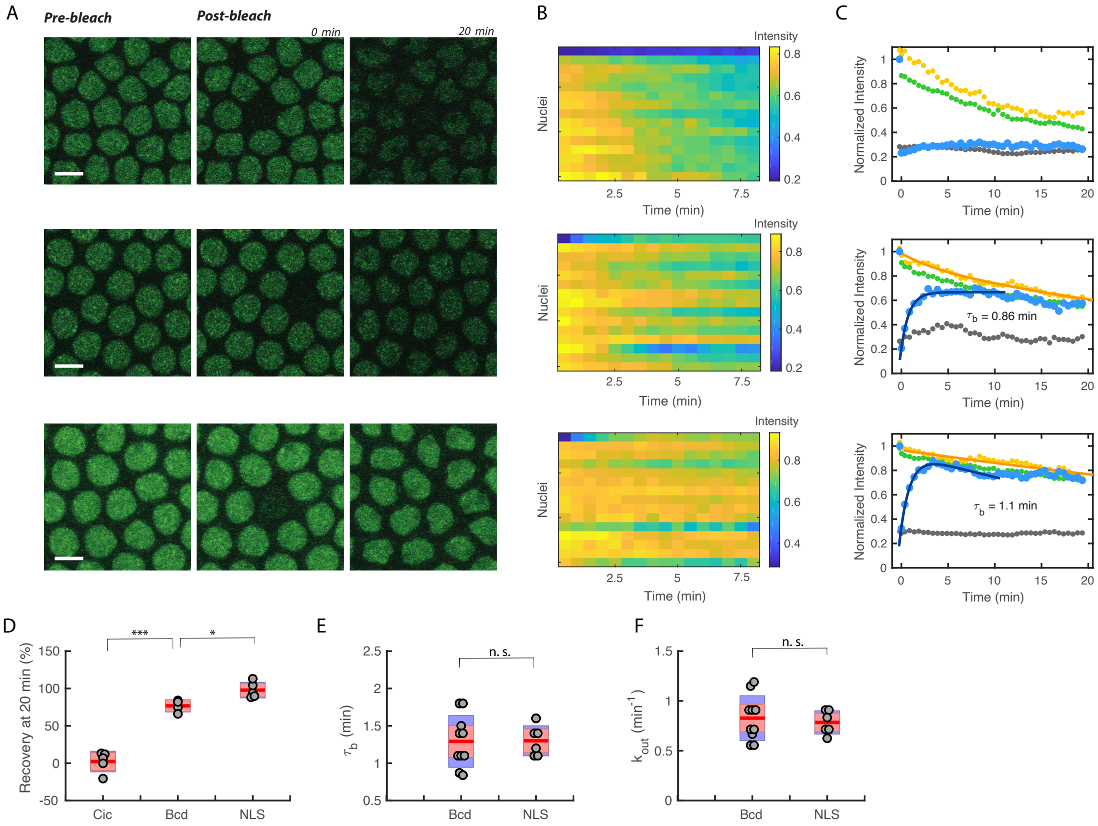


Figure S1: **Nucleo-cytoplasmic exchange of Bcd and Cic.** (**A**) Confocal images of nuclei in *D. melanogaster* embryos expressing Cic-sfGFP (top), Bcd-eGFP (middle), and NLS-eGFP (bottom) pre-bleach, immediately post-bleach (t = 0 min) and t = 20 min post-bleach. Scale bar is 5 $\mu$m. (**B**) Carpet plot showing the time evolution of the mean fluorescence intensity (normalized to the pre-photobleaching intensity) of the photobleached nucleus (top row) and 15 closest nuclei in order of proximity (top to bottom). (**C**) Mean intensity of the photobleached nucleus (blue circles), the 5 nearest-neighbour nuclei (green circles), the 5 furthest away nuclei (yellow circles) and the cytoplasm (grey circles). All intensities are normalized to the average pre-photobleaching nuclear intensity. Solid lines show fits with either a single exponential (far away nuclei, orange line) or a double exponential (photobleached nucleus, blue line, the corresponding recovery time $\tau_b$ is indicated). (**D**) Percent recovery at 20 min post-bleach measured for all three proteins. (**E**) Measured values of $\tau_b$. (**F**) Values of $k_{\text{out}} = 1/\tau_b$. In (**D-F**) the mean is indicated by the tick red line, the 95% confidence interval by the red box, and the standard deviation by the purple box. Three stars ($***$) indicate a $p$-value from the t-test less than 0.001, one star ($*$) less than 0.05, and n.s. larger than 0.5.

Table S1: Nucleo-cytoplasmic transport parameters (mean $\pm$ standard deviation) obtained from nuclear FRAP experiments.

| Experimental method | Parameters | Cic | Bcd | NLS |
|---|---|---|---|---|
| Nuclear FRAP | Recovery at 20 min | $2 \pm 14\,\%$ | $77 \pm 8\,\%$ | $98 \pm 11\,\%$ |
| | $\tau_b$ (min) | - | 1.29 (0.35) | 1.30 (0.20) |
| | $k_{\text{out}}$ (min$^{-1}$) | - | 0.83 (0.22) | 0.78 (0.12) |

### 2 Stick-and-diffuse model

#### 2.1 Description

Although only the analysis done with a two-component model is described in the paper, all the FCS data was also analyzed using a stick-and-diffuse model, which is based on the premise that diffusing fluorophores (in this case transcription factors) can transiently associate with an immobile or very slow structure (for example DNA or protein hubs bound to the DNA) therefore becoming temporarily immobilized. This model thus considers only one population, alternating between a freely diffusing state (diffusion coefficient $D$) and a transiently bound state. The part of the ACF reflecting fluorophore dynamics is then (Yeung et al., 2007):

$$
\begin{aligned}
G_D(\tau) = G_D(0) \times [&\frac{e^{-k_{\text{off}}\tau}}{1+\frac{k_{\text{off}}}{k_{\text{on}}}} + \frac{1}{1+\frac{k_{\text{on}}}{k_{\text{off}}}} \frac{e^{-k_{\text{on}}\tau}}{(1+\tau/\tau_D)\sqrt{1+\tau/(S^2\tau_D)}} \\
&+ \frac{k_{\text{on}}k_{\text{off}}}{k_{\text{on}}+k_{\text{off}}} \sum_{n=1}^{\infty} \frac{1}{(n-1)!n!} \int_0^{\tau} ds \frac{e^{-k_{\text{off}}(\tau-s)-k_{\text{on}}s}}{(1+s/\tau_D)\sqrt{1+s/(S^2\tau_D)}}(2n \\
&+ k_{\text{off}}s + k_{\text{on}}(\tau-s))(k_{\text{on}}k_{\text{off}}s(\tau-s))^{n-1}].
\end{aligned} \tag{S3}
$$

$k_{\text{on}}$ and $k_{\text{off}}$ are the binding and unbinding rate, respectively. $\tau_D$ is he characteristic diffusion time of molecules in the unbound state. It has been tested previously that $n$ can be truncated at 7 in the Taylor series (Abu-Arish et al., 2009) thus we set $n = 1, 2, ..., 7$ when fitting data with Eq. S3. The dissociation constant $K_D$ represents the equilibrium between bound and unbound species, and can be calculated through $K_D = k_{\text{on}}/k_{\text{off}}$. A quantity representing the relative amplitude of the free diffusion term can be estimated as $p = n_f/(n_f + n_b) = 1/(1 + K_D)$.

#### 2.2 Fit of FCS data

A comparison of the goodness of fit obtained when using either the stick-and-diffuse model or the two-component model for the three types of proteins studied is shown in Fig. S2. The goodness of fit is good for both models, with the two-component model resulting in a very slightly lower mean-squared difference between fit and data. For this data set, the two-component model gives a more precise estimate of $\tau_D$. The parameters characterizing a potential binding to and unbinding from DNA or hubs obtained from fitting the autocorrelation functions with the stick-and-diffuse model, namely the effective binding rate, $k_{\text{on}}$, the unbinding rate, $k_{\text{off}}$ and the fraction of unbound diffusing molecules, $p = k_{\text{off}}/(k_{\text{on}} + k_{\text{off}})$ are shown in Fig. S3.

A summary of the values of the parameters obtained from fitting the FCS data with the stick-and-diffuse model is given in Table S2. The values of $k_{\text{on}}$ and $k_{\text{off}}$ are consistent with the timescale for interactions between Bcd and DNA that have been reported previously from single particle tracking experiments (Mir et al., 2017). However, it is hard to reconcile the stick-and-diffuse model with the fact that the bound fraction of TFs then appears to decrease over time (Fig. S3G,H). Instead, since in this model each TF is supposed to alternate quickly between a free and a bound state, photobleaching should not alter the proportion of bound to free molecules.

#### 2.3 Simulations

To simulate transient binding according to a stick-and-diffuse model, particles were randomly assigned one of two states at creation ("unbound" with a probability $p$ or "bound" with a probability $p - 1$). At each step in the simulation unbound particles were allowed to diffuse with a diffusion coefficient $D_f$ and change state with a probability $k_{on}\delta t$. Bound particles, on the other hand, were not allowed to move ($D = 0$) but were allowed to change state with a probability $k_{off}\delta t = p/(1-p)k_{on}\delta t$.

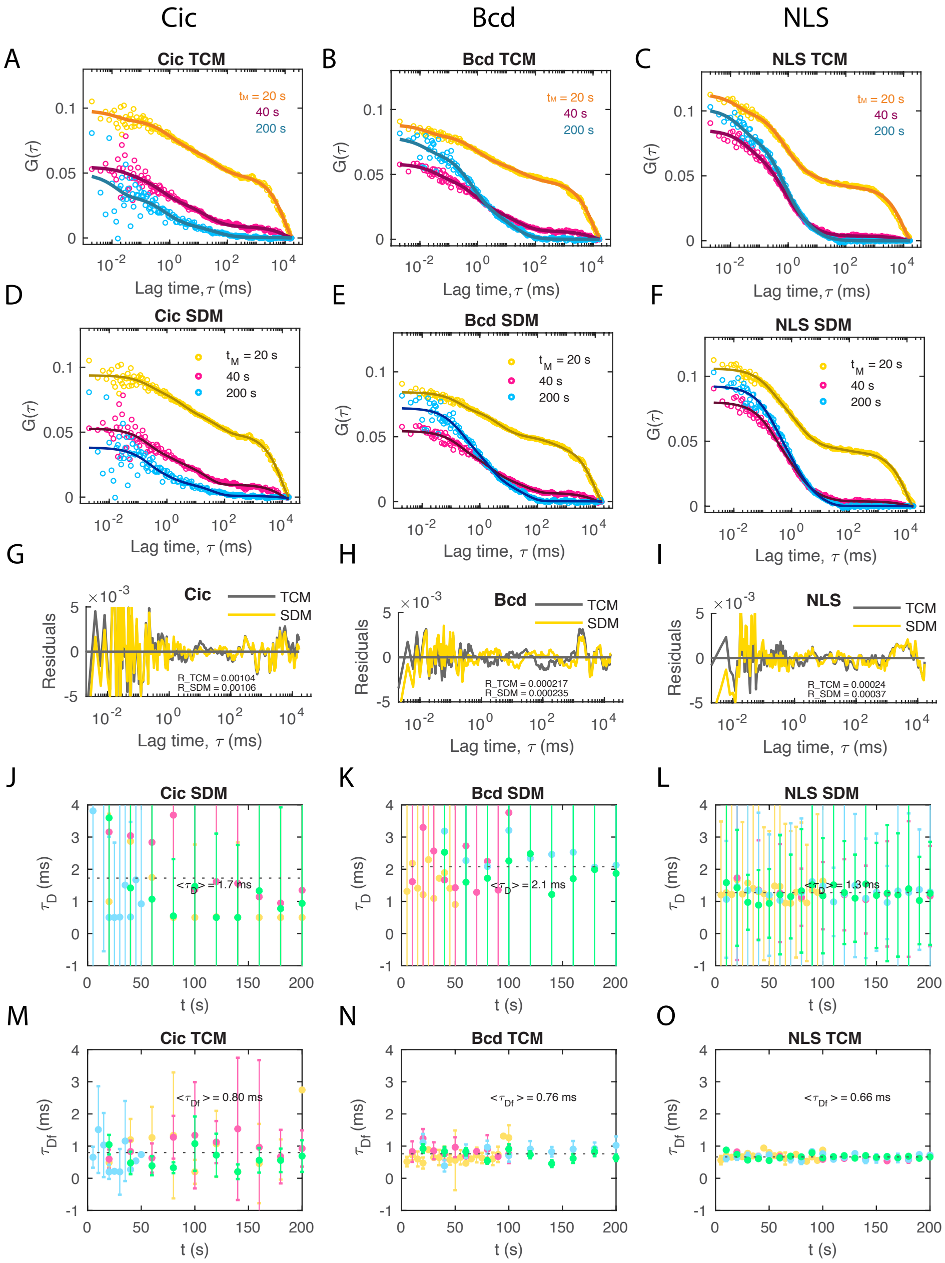


Figure S2: **Comparison between the two-component model (TCM) and the stick-and-diffuse model (SDM).** (**A,B,C**) Representative data (open circles) and fit with the TCM including a photobleaching term (solid line) of the first (orange), second (pink) and tenth (blue) ACFs from a series of single-point FCS measurements in the center of a nucleus in Drosophila embryos expressing Cic-sfGFP (**A**), Bcd-eGFP (**B**), or NLS-eGFP (**C**). (**D,E,F**) Same ACFs as in (**A,B,C**). Each measurement in the series lasted 20 s in this case, thus the first, second and tenth ACFs corresponds to $t_M = 20$ s, 40 s, and 200 s, respectively. (**G,H,I**) Comparison of the residuals obtained from fitting the first ACF with either the TCM (grey line) or the SDM (yellow line). The value of the sum of the mean-squared residuals is indicated. To further compare the goodness of fit from the two models, the characteristic time $\tau_D$ obtained from the SDM (**J,K,L**) and the fast diffusion characteristic time $\tau_{\text{Df}}$ from the TCM (**M,N,O**) are also shown, where symbols of different color represent different series of data with different measurement time (5, 10, 20s). Error bars are 50 % confidence intervals.

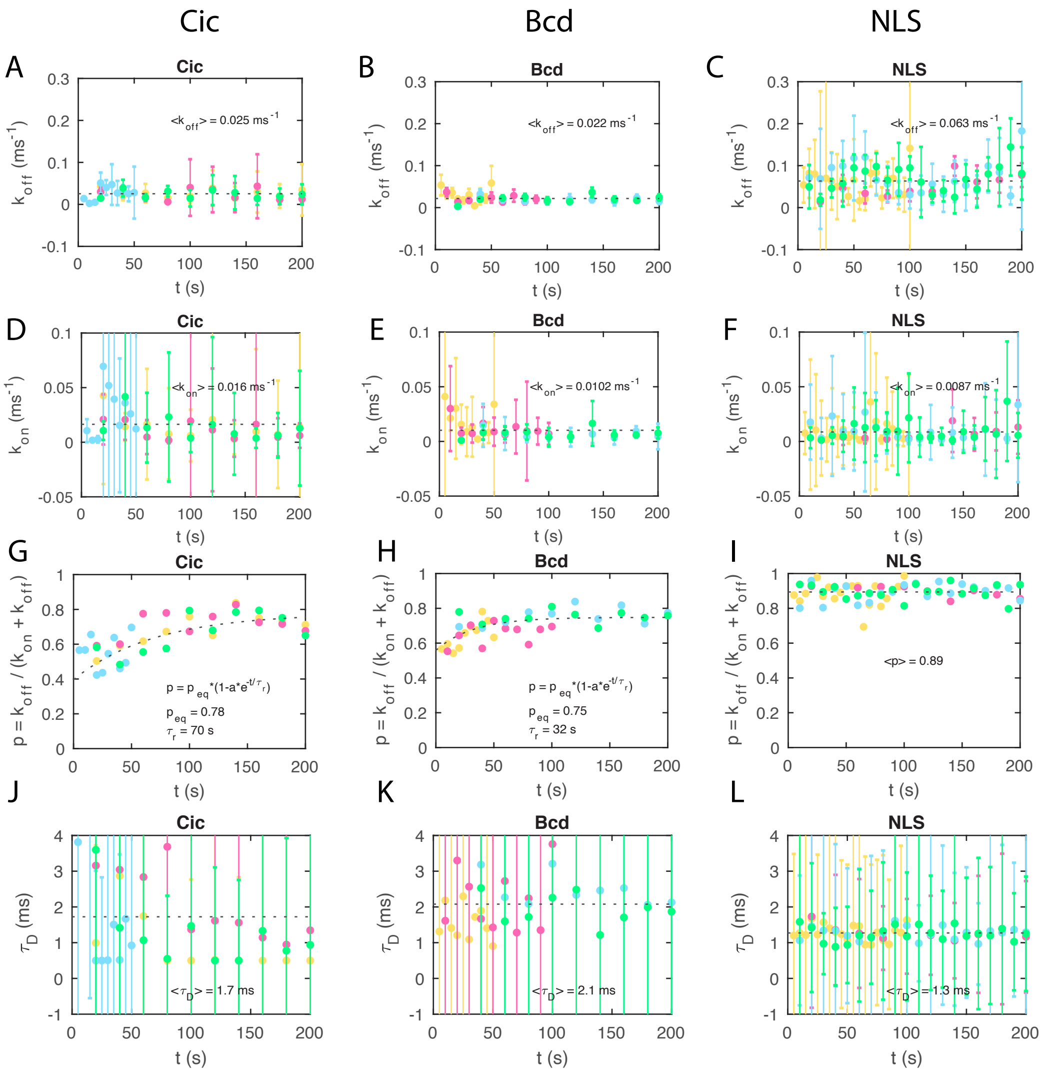


Figure S3: **Parameters extracted from fit of the data with the stick-and-diffuse model.** The unbinding rate, $k_{\text{off}}$, binding rate, $k_{\text{on}}$, the effective fast fraction, $p = k_{\text{off}}/(k_{\text{on}} + k_{\text{off}})$, and the characteristic diffusion time, $\tau_D$, from fitting the ACFs with Eq. S3, as well as their averaged values represented by dotted grey lines are shown. Different colors of symbols represents different series of measurement with measurement time of 5, 10 or 20 seconds. Error bars represent 50% confidence intervals.

Table S2: Parameters obtained using the stick-and-diffuse model to fit ACFs (mean ± standard deviation).

| Parameter | Symbol | Cic | Bcd | NLS | Unit |
|---|---|---|---|---|---|
| Binding rate | $k_{\text{on}}$ | $16 \pm 15$ | $10 \pm 9$ | $9 \pm 8$ | $s^{-1}$ |
| Unbinding rate | $k_{\text{off}}$ | $25 \pm 12$ | $22 \pm 12$ | $63 \pm 32$ | $s^{-1}$ |
| Equilibrium constant | $K_D$ | $1.6 \pm 2.2$ | $2.2 \pm 3.2$ | $7.2 \pm 10$ | |
| Diffusion coefficient | $D_f$ ($\mu m^2/s$) | $13 \pm 15$ | $11 \pm 5$ | $18 \pm 3$ | $\mu m^2/s$ |
| Fraction bound ($t = 0$) | $p_0$ | $0.38 \pm 0.06$ | $0.56 \pm 0.07$ | $0.89 \pm 0.05$ | |
| Fraction bound ($t \to \infty$) | $p_\infty$ | $0.77 \pm 0.04$ | $0.75 \pm 0.04$ | $0.89 \pm 0.05$ | |

### 3 Characteristic continuous photobleaching time

Table S3 shows the values of the characteristic continuous photobleaching time ($\tau_p$) for Cic-sfGFP, Bcd-eGFP and NLS-eGFP estimated using different methods, namely the fitting of the fluorescence count rate with an exponential function, the fitting of the ACFs with either a two-component or a stick-and-diffuse model, or the fitting of $p(t)$ (obtained either from analysis of the ACF with a two-component or a stick-and-diffuse model) with an exponential function.

Table S3: Values of $\tau_p$ obtained by different methods (mean ± standard deviation).

| Model | Method | Cic | Bcd | NLS | Unit |
|---|---|---|---|---|---|
| - | Fit of $I(t)$ | $28 \pm 6$ | $23 \pm 7$ | $22 \pm 3$ | s |
| Two-component model | Fit of $G(\tau)$ | $33 \pm 19$ | $43 \pm 24$ | $34 \pm 17$ | s |
| | Fit of $p(t)$ | $46 \pm 9$ | $37 \pm 16$ | - | s |
| Stick-and-diffuse model | Fit of $G(\tau)$ | $52 \pm 23$ | $62 \pm 34$ | $55 \pm 36$ | s |
| | Fit of $p(t)$ | $48 \pm 24$ | $32 \pm 11$ | - | s |

### 4 Determination of the localization precision

The localization precision, $\sigma^2$, of fluorescent clusters detected in confocal images, was estimated both for experimental images and simulated images, in the exact same conditions as our experiments and simulations, using control samples and control simulations.

#### 4.1 Control experimental samples

Samples of 100 nm yellow/green fluorescent beads in agar were prepared to obtain movies of immobile fluorescent beads. Sample holders were prepared by placing and melting two strips of parafilm between a microscope slide and a coverslip, so as to create a channel. A 1.5% agar solution was mixed with a diluted bead dilution and heated in a microwave until bubbling, then pipetted into a pre-warmed sample holder channel. The ends of the channel were sealed using clear nail polish. Samples were kept wrapped in foil and refrigerated until imaging. Confocal image series of the immobilized beads were captured in the exact same conditions as the live embryos, except for the laser power which was varied between 0.1% and 0.5% to achieve a range of signal-to-noise ratios (SNRs). The region of interest (ROI) was photobleached by scanning at high laser intensity for a set time in order to reduce the SNR of the beads prior to imaging. Examples of images obtained at different laser powers are shown in Fig. S4A.

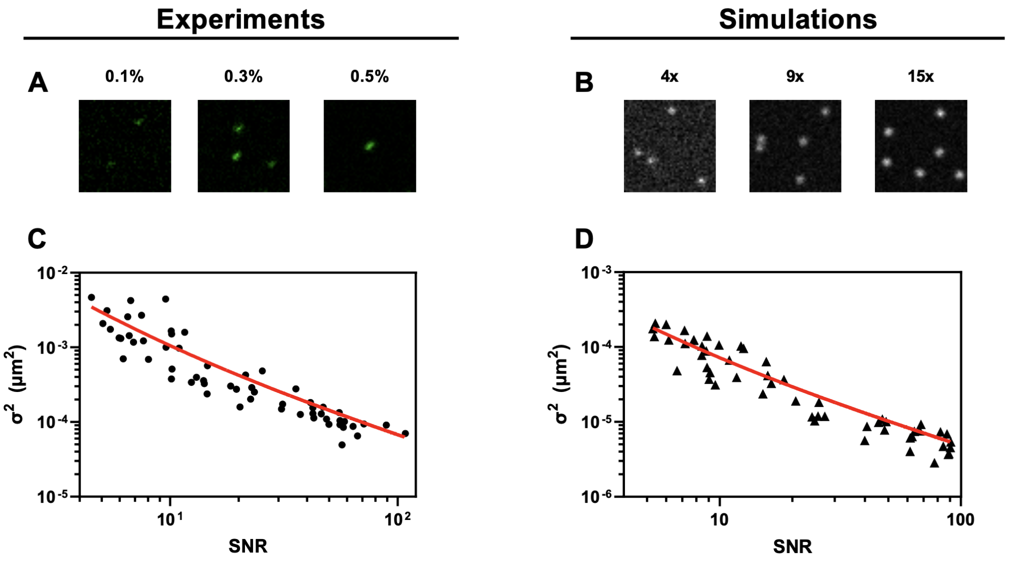


$$\langle\sigma^2\rangle = \frac{0.006(3)}{SNR} + \frac{0.04(2)}{SNR^2} \qquad \langle\sigma^2\rangle = \frac{0.0005(1)}{SNR} + \frac{0.0026(9)}{SNR^2}$$

Figure S4: **Localization precision of particle detection.** (A) Example images of beads immobilized in agar acquired with an excitation power of 0.1%, 0.3%, and 0.5% of the total laser power (2 mW). (B) Examples of simulated confocal images of immobile particles with brightness equal to 4×, 9×, and 15× that of the background in a single pixel. (C, D) Plots of the measured localization precision against the SNR for the experimental images (C) and simulated images (D). The result of the fit (red curves) are given below the plot.

##### 4.1.1 Control simulations

The localization precision for the simulated confocal images was also estimated, by simulating movies of immobile particles following the same method as outlined in the main text for generating simulated time-lapse confocal image movies. Simulations were run for particles with different brightness in order to reproduce situations with different SNRs. Examples of simulated images for different particle brightness are provided in Fig. S4B.

##### 4.1.2 Image analysis

Fluorescent particles were detected in the control movies with the FIJI mosaic plugin using the exact same parameters as for live embryos imaging and simulated images of nuclei. An apparent diffusion coefficient was obtained from the MSD of the detected beads or simulated immobile fluorophores. In parallel, the intensity of each bead was measured using the plot profile function in FIJI. The background noise was estimated as the standard deviation of the pixel intensities within a ROI containing no bead. The SNR was then calculated as the intensity of the bead divided by the background noise. The obtained localization precision ($\sigma^2 = 4D\Delta t$) is plotted as a function of SNR for the experimental immobilized beads control sample (Fig. S4C), and for the simulated control samples (Fig. S4D). In both case we expect:

$$\sigma^2 = \frac{A}{SNR} + \frac{B}{SNR^2} \tag{S4}$$

where the first term is due to photon noise and the second term is due to background noise Thompson et al. (2002). The data was fit with Eq. S4 to obtain an estimate of the localization precision as a function of SNR for our confocal imaging system and for our simulations.
